# Struo2: efficient metagenome profiling database construction for ever-expanding microbial genome datasets

**DOI:** 10.1101/2021.02.10.430604

**Authors:** Nicholas D. Youngblut, Ruth E. Ley

## Abstract

Mapping metagenome reads to reference databases is the standard approach for assessing microbial taxonomic and functional diversity from metagenomic data. However, public reference databases often lack recently generated genomic data such as metagenome-assembled genomes (MAGs), which can limit the sensitivity of read-mapping approaches. We previously developed the Struo pipeline in order to provide a straight-forward method for constructing custom databases; however, the pipeline does not scale well with the ever-increasing number of publicly available microbial genomes. Moreover, the pipeline does not allow for efficient database updating as new data are generated. To address these issues, we developed Struo2, which is >3.5-fold faster than Struo at database generation and can also efficiently update existing databases. We also provide custom Kraken2, Bracken, and HUMAnN3 databases that can be easily updated with new genomes and/or individual gene sequences. Struo2 enables feasible database generation for continually increasing large-scale genomic datasets.

**Availability:**

- Struo2: https://github.com/leylabmpi/Struo2
- Pre-built databases: http://ftp.tue.mpg.de/ebio/projects/struo2/
- Utility tools: https://github.com/nick-youngblut/gtdb_to_taxdump

## Results

Metagenome profiling involves mapping reads to reference sequence databases and is the standard approach for assessing microbial community taxonomic and functional composition via metagenomic sequencing. Most metagenome profiling software includes “standard” reference databases. For instance, the popular HUMANnN pipeline includes multiple databases for assessing both taxonomy and function from read data (Franzosa *et al.*, 2018). Similarly, Kraken2 includes a set of standard databases for taxonomic classification of specific clades (*e.g.,* fungi or plants) or all taxa (Wood *et al.*, 2019). While such standard reference databases provide a crucial resource for metagenomic data analysis, they may not be optimal for the needs of researchers. For example, a custom database that includes newly generated MAGs can increase the percent of reads mapped to references (Youngblut *et al.*, 2020). The process of making custom reference databases is often complicated and requires substantial computational resources, which led us to create Struo for straight-forward custom metagenome profiling database generation (de la Cuesta-Zuluaga *et al.*, 2020). However, Struo requires ~2.4 CPU hours per genome, which would necessitate >77,900 CPU hours (>9.1 years) if including one genome per the 31,911 species in Release 95 of the Genome Taxonomy Database (GTDB) (Parks *et al.*, 2018).

Struo2 generates Kraken2 and Bracken databases similarly to Struo (Lu *et al.*, 2017; Wood *et al.*, 2019), but the algorithms diverge substantially for the time consuming step of gene annotation required for HUMAnN database construction. Struo2 performs gene annotation by clustering all gene sequences of all genomes using the *mmseqs2 linclust* algorithm, and then each gene cluster representative is annotated via *mmseq2 search* (Figure 1A; Supplemental Methods) (Steinegger and Söding, 2017, 2018). In contrast, Struo annotates all non-redundant genes of each genome with DIAMOND (Buchfink *et al.*, 2015). Struo2 utilizes snakemake and conda, which allows for easy installation of all dependencies and simplified scaling to high performance computing systems (Köster and Rahmann, 2012).

**Figure 1.**
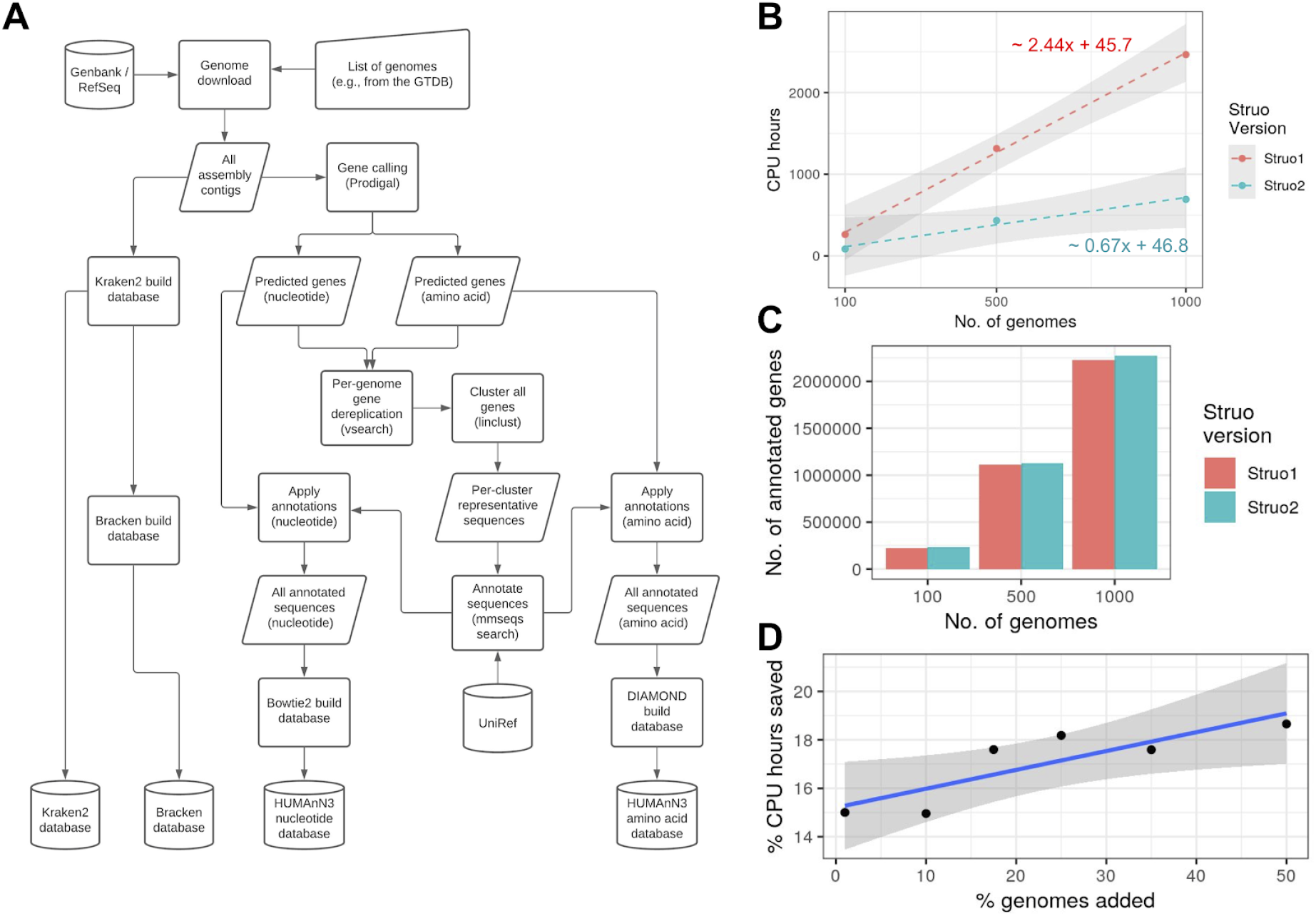
Struo2 can build databases faster than Struo and can efficiently update the databases. A) A general outline of the Struo2 database creation algorithm. Cylinders are input or output files, squares are processes, and right-tilted rhomboids are intermediate files. The largest change from Struo is the utilization of mmseqs2 for clustering and annotation of genes. B) Benchmarking the amount of CPU hours required for Struo and Struo2, depending on the number of input genomes. C) The number of genes annotated with a UniRef90 identifier. D) The percent of CPU hours saved via the Struo2 database updating algorithm versus *de novo* database generation. The original database was constructed from 1000 genomes. For B) and D), the grey regions represent 95% confidence intervals.

Benchmarking on genome subsets from the GTDB showed that Struo2 requires ~0.67 CPU hours per genome versus ~2.4 for Struo (Figure 1B). Notably, Struo2 annotates slightly more genes than Struo, possibly due to the sensitivity of the *mmseqs search* iterative search algorithm (Figure 1C). The use of mmseqs2 allows for efficient database updating of new genomes and/or individual gene sequences via *mmseqs clusterupdate* (Figure S1); we show that this approach saves 15-19% of the CPU hours relative to generating a database from scratch (Figure 1D).

We used Struo2 to create publicly available Kraken2, Bracken, and HUMAnN3 custom databases from Release 95 of the GTDB (see Supplemental Methods). We will continue to publish these custom databases as new GTDB versions are released. The databases are available at http://ftp.tue.mpg.de/ebio/projects/struo2/. We also created a set of utility tools for generating NCBI taxdump files from the GTDB taxonomy and mapping between the NCBI and GTDB taxonomies. The taxdump files are utilized by Struo2, but these tools can be used more generally to integrate the GTDB taxonomy into existing pipelines designed for the NCBI taxonomy (available at https://github.com/nick-youngblut/gtdb_to_taxdump).

## Data availability

Struo2 is available at https://github.com/leylabmpi/Struo2, the pre-built databases can be found at http://ftp.tue.mpg.de/ebio/projects/struo2/, and utility tools are located at https://github.com/nick-youngblut/gtdb_to_taxdump.

## Acknowledgements

This study was supported by the Max Planck Society. We thank Albane Ruaud, Liam Fitzstevens, Jacobo de la Cuesta-Zuluaga, and Jillian Waters for providing helpful comments on an earlier version of this manuscript.

