## Supplemental Materials for "Struo2: efficient metagenome profiling database construction for ever-expanding microbial genome datasets"

**Supplemental Methods**

*Struo2 database creation algorithm*

Struo2 can generate database files for 4 main database types: “Kraken2”, “Bracken”, “genes”, and “HUMAnN3” (Wood *et al.*, 2019; Lu *et al.*, 2017; Franzosa *et al.*, 2018). Struo2 uses snakemake and conda (Köster and Rahmann, 2012), and so there are no dependencies that must be installed prior to pipeline execution besides snakemake, conda, and pandas (for input table loading). Moreover, snakemake allows for efficient job execution and easy scaling on to high performance computing systems. We note that the Struo2 pipeline code is a substantial re-write and expansion of the original Struo pipeline (*e.g.,* ~1500 versus ~7000 lines of code in Struo versus Struo2, respectively).

The user input for Struo2 database creation is a table that lists: i) unique taxon names, ii) assembly accession identifiers (if available), iii) paths to (compressed) genome assembly fasta files, iv) taxonomy identifiers (taxids) used for Kraken2 database construction, and v) taxonomies at the genus and species levels (used for HUMAnN3). We provide 2 utility scripts to aid in construction of custom databases from genomes in the GTDB: *GTDB_metadata_filter.R* and *genome_download.R*. *GTDB_metadata_filter.R* can filter the publicly available GTDB archaeal and bacterial genome metadata files to a select subset of genomes (*e.g.,* those with a lower CheckM-estimated contamination). *genome_download.R* can then download all of the user-selected GTDB genomes and add the path to the genome assembly fasta files to the GTDB metadata table. This updated metadata table can then be directly used as input to GTDB.

For construction of the custom Kraken2 database, contigs are renamed to “kraken:taxid|<taxid>|<seqid>”, as described in the Kraken2 manual (https://github.com/DerrickWood/kraken2/wiki/Manual). The renamed contigs are added to a new Kraken2 database via *kraken-build*, and then the database is constructed via the same command. By default, the GTDB taxonomy is used, which entails providing custom GTDB taxdump files created via the *gtdb_to_taxdump.py* utility tool (available at https://github.com/nick-youngblut/gtdb_to_taxdump). The “taxonomy” and “library” directories created by Kraken2 for temporary file storage are saved in order to expedite database updating with new genomes.

Custom Bracken database files are created for any number of read lengths that the user specifies (100 and 150 base pairs by default). The *bracken-build.py* script is used within the pipeline for constructing each Bracken database.

In order to construct a custom HUMAnN3 database, Struo2 first creates a precursor “genes” database, which consists of gene sequences from each genome and gene clusters generated via *mmseqs linclust*. To construct the “genes” database, genes are first called via prodigal (Hyatt *et al.*, 2010), and then de-replicated at 97% sequence identity with vsearch (Rognes *et al.*, 2016), which is similar to the standard HUMAnN database construction process (Franzosa *et al.*, 2018). Non-redundant gene sequences from all genomes are combined, and the metadata of each gene sequence (*e.g.,* genome of origin, contig of origin, and location on the contig) is also combined into one text file. The amino acid gene sequences are clustered via *mmseqs linclust*. By default, gene cluster representative sequences are annotated against UniRef90 (version 2019-01; the same as used by HUMAnN3) via *mmseqs search* with 2 search iterations and 3 sensitivity steps (min=1, max=6). Prior to annotation, the sequence queries are split into *n* batches and run in parallel for faster distributed searching with snakemake (*n* is user-defined). For each gene cluster, the UniRef90 annotations are propagated to each gene. UniRef90 annotations are mapped to UniRef50 identifiers via a mapping file created from the UniRef90.xml file available from the UniProt ftp server (ftp://ftp.uniprot.org/pub/databases/uniprot/). The *unirefxml2clust50-90idx.py* utility script is used to generate this mapping file (available at https://github.com/nick-youngblut/gtdb_to_taxdump). The mapping of UniRef90 to UniRef50 identifiers obviates the need to annotate genes separately against UniRef90 and UniRef50. We note that Struo requires separate rounds of annotation to each UniRef database instead of this UniRef90-to-UniRef50 mapping approach, which greatly increases the run time versus Struo2 when the goal is to obtain annotations for both UniRef90 and UniRef50. Note that the genes database includes both nucleotide and amino acid sequences for each gene.

The annotated gene sequences are renamed in the format "<UniRefID>|<gene_length>|g__<genus>;s__<species>" for creation of the HUMAnN3 database. Note that the taxonomy information is provided by the user in the original input table. *bowtie2-build* and *diamond makedb* are used to generate a HUMAnN3-compatible bowtie2 and DIAMOND databases of all annotated gene nucleotide and amino acid sequences, respectively.

*Struo2 database update algorithm*

Struo2 can update existing Struo2-generated Kraken2, Bracken, genes, and HUMAnN3 databases. The databases can be updated with new genomes or individual gene sequences (*e.g.,* created via metagenome assembly with PLASS (Steinegger *et al.*, 2019)).

If the input is a set of new genomes, the input is essentially the same as for database creation, except the existing database files must also be provided. Database updating with individual gene sequences requires the gene sequences in amino acid format (and also nucleotide, if available) and metadata on each gene (*i.e.,* the genus- and species-level taxonomy inferred via *mmseqs taxonomy* or other approaches).

Kraken2 custom databases are updated via adding more genomes to the existing library via *kraken-build*. New Bracken databases are created from the updated Kraken2 database.

Gene sequences, either originating from new genomes or new individual sequences, are added to the existing mmseqs gene cluster database via *mmseqs clusterupdate*. Newly formed clusters are annotated with *mmseqs search*, while existing annotations are used for existing clusters. The updated database of annotated genes are used for creating new HUMAnN3-compatible bowtie2 and DIAMOND databases.

*Benchmarking*

We used genomes from the GTDB (Release 95) for all benchmarking. Only genomes with ≥50% CheckM-estimated completeness, <5% CheckM-estimated contamination were included (Parks *et al.*, 2015). To reduce biases towards species with large numbers of representative genomes, we selected one genome per species. The genome with the highest estimated completeness and lowest estimated contamination was selected for all candidates of each species. The final pool consisted of 30,989 genomes (Figure S2).

We used the same genome subsets for benchmarking database creation with both Struo and Struo2. We benchmarked the combined time to generate Kraken2, Bracken, and HUMAnN databases, which included both UniRef50 and UniRef90 annotations for the HUMAnN databases. Both pipelines were run on the same computational architecture, consisting of a high performance computing cluster comprising nodes running Ubuntu 18.04.5 with AMD Epyc CPUs and 0.5-2 terabytes of RAM. The CPU hours shown in Figure 1B are the sum of all CPU hours for all snakemake jobs, as recorded via snakemake's benchmarking feature.

We only benchmarked database updating for Struo2, given that Struo cannot update databases, and we clearly show in Figure 1B that database generation is much slower for Struo. We first used Struo2 to generate custom Kraken2, Bracken, and HUMAnN databases from 1000 genomes. These "n1000" databases were used for all database update benchmarking. The genomes used for database update benchmarking did not overlap with any genomes used to generate the n1000 databases, and they did not overlap with each other. We used subsets of 10, 100, 175, 250, 350, and 500 genomes. We used the linear regression models shown in Figure 1B to estimate the CPU hours that would be required to generate each database from scratch rather than updating.

*Struo2 databases from GTDB Release 95*

The genomes selected were as reported for the benchmarking of Struo and Struo2. The custom Kraken2, Bracken, genes, and HUMAnN3 databases are available at: http://ftp.tue.mpg.de/ebio/projects/struo2/. We will publish new versions of each database as new releases of the GTDB are published.

*Utility tools*

We have generated a set of utility tools for aiding in the construction of input for Struo2 and generally facilitating the integration of the GTDB taxonomy into existing bioinformatics pipelines. Some of these tools are described elsewhere in the Supplement Methods. We note 2 utility tools that can have a broad applicability: *gtdb_to_taxdump.py* and *ncbi-gtdb_map.py*. The former can convert the GTDB taxonomy, as documented in the GTDB bacterial and archaeal metadata table, to NCBI-formatted taxdump files. These taxdump files can be used with any existing software that requires taxdump files, such as taxonkit (Shen and Xiong, 2019) or KrakenUniq (Breitwieser *et al.*, 2018). *ncbi-gtdb_map.py* maps between NCBI and GTDB taxonomies, based on the taxonomy information provided in the GTDB archaeal and bacterial metadata files. This tool can be useful for converting GTDB-Tk classifications to NCBI taxonomies (Chaumeil *et al.*, 2019), or converting existing NCBI taxonomies to GTDB taxonomies without requiring re-classification.

### Supplemental Figures

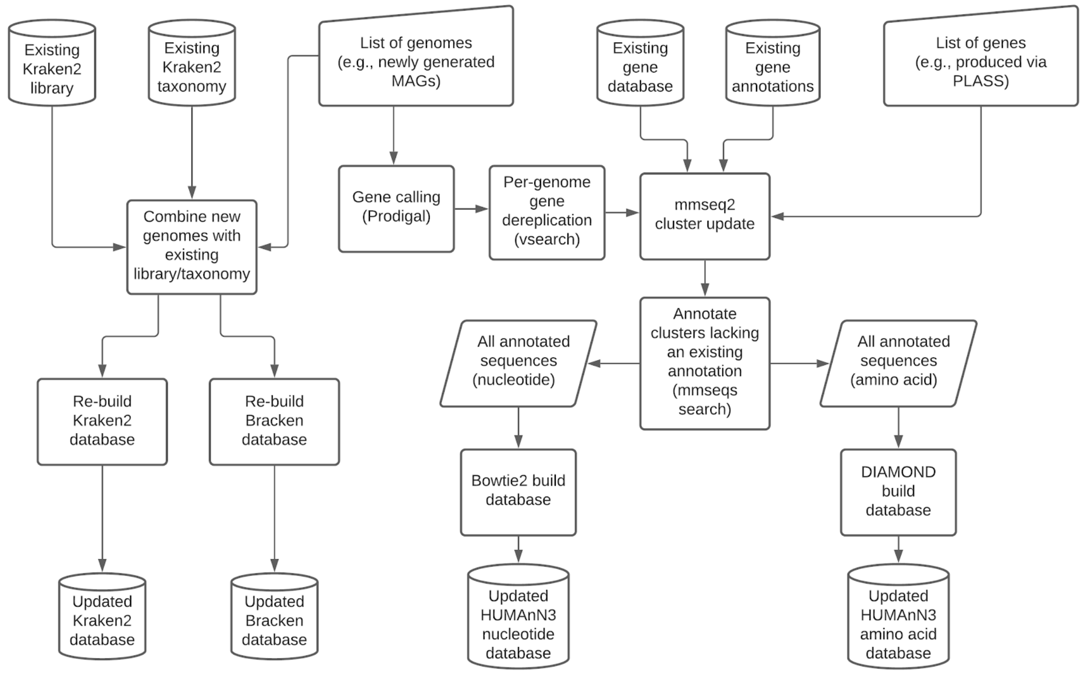


**Figure S1.** An overview of the Struo2 algorithm for database updating. Cylinders are input or output files, squares are processes, and right-tilted rhomboids are intermediate files. Existing Kraken2, Bracken, genes, and HUMAnN3 databases can be updated with new genomes, while only existing genes and HUMAnN3 databases can be updated with new individual gene sequences.

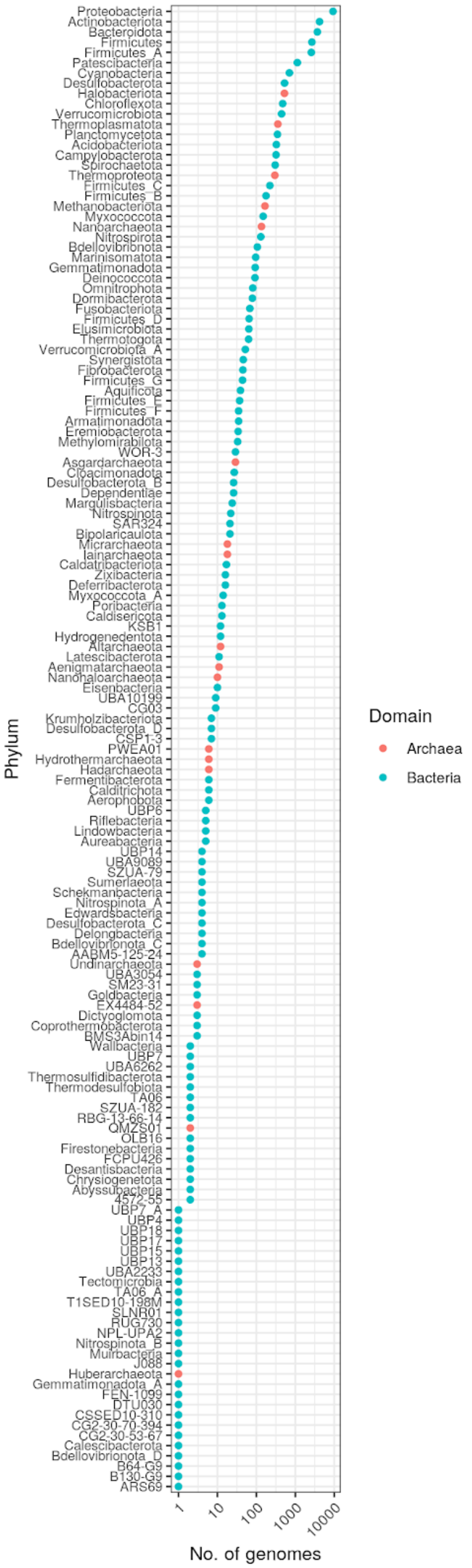
Phylum
No. of genomes
1
10
100
1000
10000
Domain
Archaea
Bacteria
Proteobacteria
Actinobacteriota
Bacteroidota
Firmicutes
Firmicutes_A
Patescibacteria
Cyanobacteria
Desulfobacterota
Halobacteriota
Chloroflexota
Verrucomicrobiota
Thermoplasmatota
Planctomycetota
Acidobacteriota
Campylobacterota
Spirochaetota
Thermoproteota
Firmicutes_C
Firmicutes_B
Methanobacteriota
Myxococcota
Nanoarchaeota
Nitrospirota
Bdellovibrionota
Marinisomatota
Gemmatimonadota
Deinococcota
Omnitrophota
Dormibacterota
Fusobacteriota
Firmicutes_D
Elusimicrobiota
Thermotogota
Verrucomicrobiota_A
Synergistota
Fibrobacterota
Firmicutes_G
Aquificota
Firmicutes_E
Firmicutes_F
Armatimonadota
Eremiobacterota
Methylomirabilota
WOR-3
Asgardarchaeota
Cloacimonadota
Desulfobacterota_B
Dependentiae
Margulisbacteria
Nitrospinota
SAR324
Bipolaricaulota
Micrarchaeota
Iainarchaeota
Caldatribacteriota
Zixibacteria
Deferribacterota
Myxococcota_A
Poribacteria
Caldisericota
KSB1
Hydrogenedentota
Altarchaeota
Latescibacterota
Aenigmatarchaeota
Nanohaloarchaeota
Eisenbacteria
UBA10199
CG03
Krumholzibacteriota
Desulfobacterota_D
CSP1-3
PWEA01
Hydrothermarchaeota
Hadarchaeota
Fermentibacterota
Calditrichota
Aerophobota
UBP6
Riflebacteria
Lindowbacteria
Aureabacteria
UBP14
UBA9089
SZUA-79
Sumerlaeota
Schekmanbacteria
Nitrospinota_A
Edwardsbacteria
Desulfobacterota_C
Delongbacteria
Bdellovibrionota_C
AABM5-125-24
Undinarchaeota
UBA3054
SM23-31
Goldbacteria
EX4484-52
Dictyoglomota
Coprothermobacterota
BMS3Abin14
Wallbacteria
UBP7
UBA6262
Thermosulfidibacterota
Thermodesulfobiota
TA06
SZUA-182
RBG-13-66-14
QMZS01
OLB16
Firestonebacteria
FCPU426
Desantisbacteria
Chrysiogenetota
Abyssubacteria
4572-55
UBP7_A
UBP4
UBP18
UBP17
UBP15
UBP13
UBA2233
Tectomicrobia
TA06_A
T1SED10-198M
SLNR01
RUG730
NPL-UPA2
Nitrospinota_B
Muirbacteria
J088
Huberarchaeota
Gemmatimonadota_A
FEN-1099
DTU030
CSSED10-310
CG2-30-70-394
CG2-30-53-67
Calescibacterota
Bdellovibrionota_D
B64-G9
B130-G9
ARS69

**Figure S2.** The number of GTDB genomes per phylum used for Struo2 generation of the custom Kraken2, Bracken, genes, and HUMAnN3 databases available at http://ftp.tue.mpg.de/ebio/projects/struo2/. See the Supplemental Methods for information on how genomes were selected. The phylum names shown are based on the GTDB taxonomy.
